# Cell lineage specification during development of the anterior lateral plate mesoderm and forelimb field

**DOI:** 10.1101/2022.01.13.475748

**Authors:** Axel H Newton, Sarah M Williams, Andrew T Major, Craig A Smith

**Affiliations:** Biomedicine Discovery Institute, Monash University, Victoria, Australia; School of BioSciences, The University of Melbourne, Victoria, Australia; Monash Bioinformatics Platform, Monash University, Victoria, Australia; QCIF, University of Queensland, Queensland, Australia

## Abstract

The lateral plate mesoderm (LPM) is a transient embryonic tissue that gives rise to a diverse range of mature cell types, including the cardiovascular system, the urogenital system, endoskeleton of the limbs, and mesenchyme of the gut. While the genetic processes that drive development of these tissues are well defined, the early cell fate choices underlying LPM development and specification are poorly understood. In this study, we utilize single-cell transcriptomics to define cell lineage specification during development of the anterior LPM and the forelimb field in the chicken embryo. We identify the molecular pathways directing differentiation of the aLPM towards a somatic or splanchnic cell fate, and subsequent emergence of the forelimb mesenchyme. We establish the first transcriptional atlas of progenitor, transitional and mature cell types throughout the early forelimb field and uncover the global signalling pathways which are active during LPM differentiation and forelimb initiation. Specification of the somatic and splanchnic LPM from undifferentiated mesoderm utilizes distinct signalling pathways and involves shared repression of early mesodermal markers, followed by activation of lineage-specific gene modules. We identify rapid activation of the transcription factor *TWIST1* in the somatic LPM preceding activation of known limb initiation genes, such as *TBX*5, which plays a likely role in epithelial-to-mesenchyme transition of the limb bud mesenchyme. Furthermore, development of the somatic LPM and limb is dependent on ectodermal BMP signalling, where BMP antagonism reduces expression of key somatic LPM and limb genes to inhibit formation of the limb bud mesenchyme. Together, these findings provide new insights into molecular mechanisms that drive fate cell choices during specification of the aLPM and forelimb initiation.

## Introduction

The lateral plate mesoderm (LPM) is a transient, embryonic tissue in vertebrate embryos which produces a remarkable diversity of cell and organ types including the cardiovascular system, urogenital system, smooth muscle and connective tissues of the limb (Nishimoto and Logan, 2016; Prummel et al., 2019, 2020). The LPM arises in the vertebrate embryo from the mesodermal germ layer as bilateral sheets along the anterior–posterior (A–P) axis, forming anterior (aLPM) and posterior (pLPM) domains (Tanaka, 2016). Diversification of the primitive mesoderm into lateral plate, paraxial (somite) or axial (notochord) mesoderm occurs in response to differential combinations of BMP, FGF or WNT signals (Tonegawa et al., 1997; Loh et al., 2016). Specifically, LPM formation is achieved through localized BMP4 signalling and antagonism of WNT signals, while paraxial mesoderm forms through WNT signals and BMP antagonism via NOGGIN (Tonegawa et al., 1997; Tonegawa and Takahashi, 1998; Yoshino et al., 2016). The primitive LPM undergoes further dorsoventral subdivision into two distinct layers separated by the embryonic coelom: the somatic LPM, which fuses with the ectoderm to form the somatopleure, and splanchnic LPM which fuses with the endoderm to form the splanchnopleure (Funayama et al., 1999). The somatic LPM gives rise to the body wall, cardiovascular system, smooth muscle, amnion and limbs, while the splanchnic LPM generates mesenteries and connective tissue lining the gut and respiratory systems (Prummel et al., 2020).

Specification of the LPM progenitors towards its diverse tissue fates are well defined and achieved through activation of key transcriptional regulators. For example, activation of the transcription factors *NKX2-5*, *TBX5* or *FOXF1,* in cardiac, somatic or splanchnic LPM cells initiate development of the heart, forelimb or gut, respectively (Mahlapuu et al., 2001; Harvey et al., 2002; Agarwal et al., 2003). However, the mechanisms that drive LPM formation and subdivision are not well defined (Prummel et al., 2019, 2020). Initial LPM specification from the primitive mesoderm is accompanied by activation of transcription factors *FOXF1*, *HAND1, OSR1* and *PRRX1* (Kuratani et al., 1994; Peterson et al., 1997; Loh et al., 2016). During LPM development and differentiation, *OSR1* becomes restricted to the coelomic epithelium / intermediate mesoderm, *PRRX1* and activation of *IRX3* to the somatic LPM, and *HAND1* and *FOXF1* to the splanchnic LPM (Funayama et al., 1999; Mahlapuu et al., 2001). Intriguingly however, *Prrx1*^−/−^, *Irx3*^−/−^ and *Osr1*^−/−^ mouse mutants do not possess an aberrant LPM phenotype, though display later defects in the limbs (Martin et al., 1995; Wang et al., 2005; Li et al., 2014). Conversely, *Hand1*^−/−^ mutants are embryonic lethal with broad developmental defects (Firulli et al., 1998) and *Foxf1*^−/−^ mutants show partial to incomplete subdivision of the LPM, expression of somatic LPM genes in the splanchnic LPM, and gut defects, suggesting a failure of splanchnic LPM commitment (Mahlapuu et al., 2001). Thus, while *FOXF1* and *HAND1* play important roles in splanchnic LPM development, those that promote somatic LPM differentiation remain unclear.

LPM subdivision and somatic LPM identity is suggested to occur through secreted BMPs from the overlying ectoderm (Funayama et al., 1999; Mahlapuu et al., 2001). In the chicken embryo, BMP2 is sufficient to activate *PRRX1* expression in the LPM, while BMP antagonism by *NOGGIN* represses activation of *PRRX1* and *IRX3* (Funayama et al., 1999; Ocaña et al., 2012). The ectodermal origin of these signals are observed where *ex vivo* culture of LPM with ectoderm initiates *PRRX1* and *IRX3* expression, but not when LPM is cultured alone (Funayama et al., 1999). Together, these observations suggest that ectodermal BMP signals activate somatic LPM genes, restrict splanchnic LPM genes and drive subdivision of the LPM. However, this hypothesis has not been directly tested. Furthermore, the specific ligands, receptors and gene expression dynamics underlying this subdivision are yet to be defined.

After LPM subdivision, forelimb development is initiated in the somatopleure through activation of the T-box transcription factor *TBX5* (Logan et al., 1998; Agarwal et al., 2003; Rallis et al., 2003). This field is defined by nested *Hox* gene expression and RA signalling (Nishimoto et al., 2015; Tanaka, 2016). Limb outgrowth begins with a localized epithelial to mesenchymal transition (EMT) of the somatic LPM, which has been proposed to be a TBX5 and FGF10 dependant process (Gros and Tabin, 2014). *TBX5* induces the activation of *FGF10*, which establishes a positive feedback loop with *FGF8* in the ectoderm to drive outgrowth of the limb bud mesenchyme (Ohuchi et al., 1997; Moon and Capecchi, 2000; Nishimoto et al., 2015). This activates a network of patterning factors and morphogens to further promote outgrowth maintenance and patterning (for comprehensive reviews see Tickle, 2015; Zuniga, 2015). However, the events immediately preceding the *TBX5*-dependant limb regulatory pathway in the somatic LPM are less well understood.

Single-cell transcriptomics have provided high-resolution analyses of cell lineage trajectories underlying multiple aspects of mesoderm and limb development (Loh et al., 2016; Scialdone et al., 2016; Gerber et al., 2018; Feregrino et al., 2019; Pijuan-Sala et al., 2019; Han et al., 2020; Johnson et al., 2020; Mahadevaiah et al., 2020). Importantly however, details regarding the cellular decisions that underlie LPM differentiation, subdivision, and commitment to a limb fate remain undetermined. In this study, we resolve the early cell fate decisions underlying differentiation of the aLPM in the developing chicken forelimb field using single cell RNA sequencing. We define the signalling pathways and ligand-receptor pairs which communicate between the germ layers and their tissue derivatives, reconstruct lineages and gene expression dynamics during LPM development, and identify likely candidates underlying EMT and initiation of the forelimb mesenchyme. Our data corroborate known interactions within the mesoderm, but also reveal novel tissue-specific markers and gene networks activated during specification of the LPM into somatic and splanchnic tissues. Notably, we identify *TWIST1* as an early marker of somatic LPM development with a likely role underlying EMT of the limb bud mesenchyme. Finally, we highlight the importance of BMP signalling underlying development of the LPM and limb bud mesenchyme. Together, these findings provide a robust overview of the developmental landscape underlying formation and specification of the aLPM.

## Results

### Transcriptional clustering of cell populations in the presumptive chicken forelimb field

To study cell fate decisions underlying development of the chicken aLPM and forelimb, we performed single-cell RNA sequencing of cell populations and generated cell-type specific clusters. Briefly, tissues corresponding to the presumptive chicken forelimb field, lateral to somites 20-25, were dissected from embryonic day (E) 1.5, 2.5 and 3.5 chicken embryos, corresponding to approximate Hamburger-Hamilton stage (HH) 10, 14 and 18, respectively (Hamburger and Hamilton, 1951) (Figure 1a). These stages cover initial development of the aLPM, subdivision of the LPM into the somatic and splanchnic layers, and limb initiation and early outgrowth (Newton and Smith, 2020). Dissected tissues were dissociated to single cells, FACS sorted to remove dead and dying cells, then processed through the 10x Chromium system. After quality filtering, a total of 15355 cells, corresponding to 5273 cells from E1.5, 6856 from E2.5 and 3226 from E3.5 embryos, were sequenced across 16779 genes. To remove low read count or low diversity cells, we applied an additional strict filtering threshold of 2000 UMI counts per cell, yielding a total of 3262 cells, with 1210 from E1.5, 1313 from E2.5 and 739 from E3.5 embryos. Cell transcriptomic relationships were visualized with global t-distributed Stochastic Neighbour Embedding (tSNE) dimension reduction, which showed a distinct separation of cell types according to cell cycle phase and embryonic stage (Figure 1b, c).

**Figure 1.**
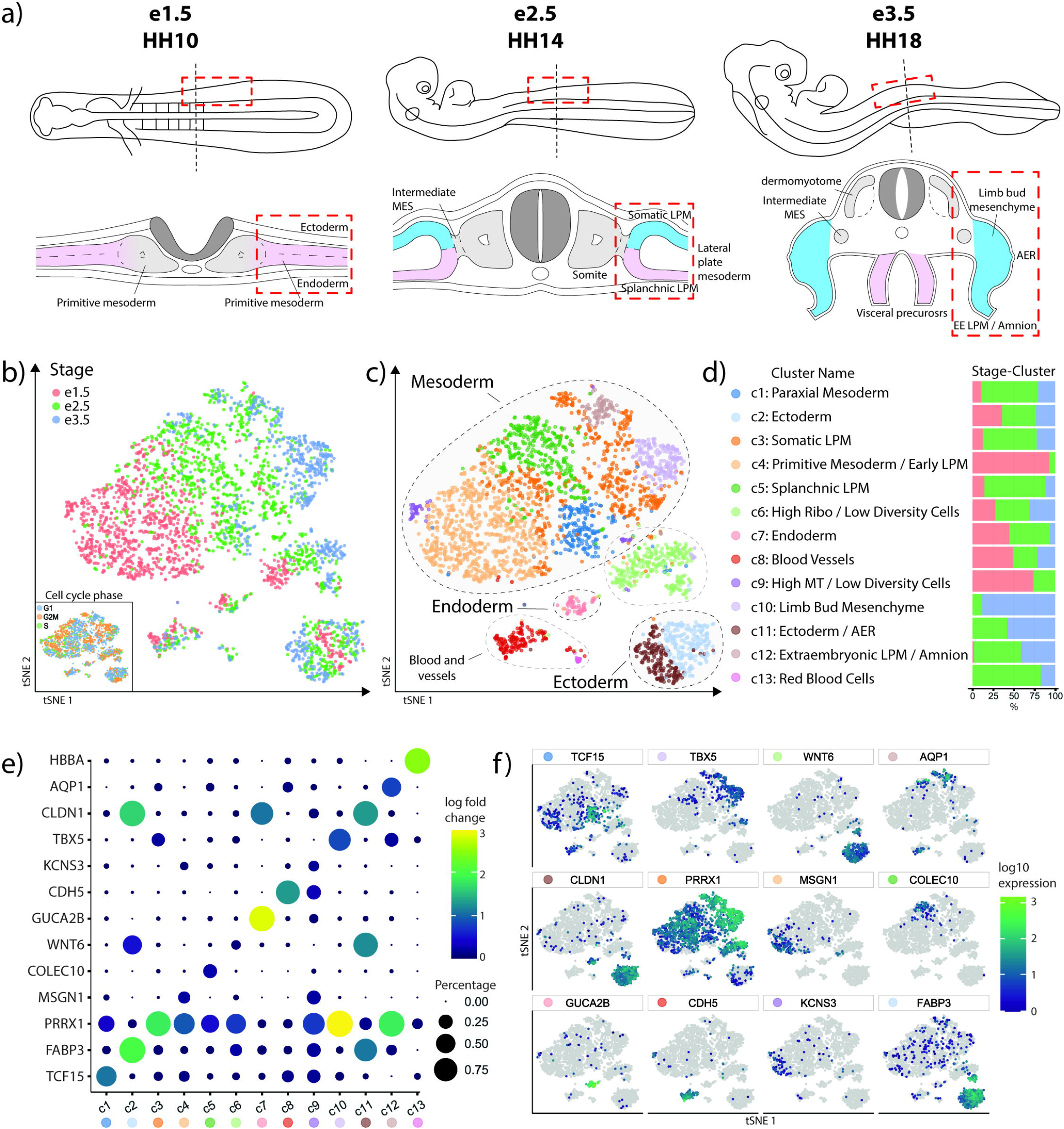
Identification of cell types in the avian forelimb field. (**A**) Cells were isolated from chicken embryonic day (e) 1.5, e2.5 and e3.5 to sample all major tissues in the developing forelimb field. (**B**) tSNE visualisation separated cells based on stage, and (**C**) germ layer origin. (**D**) Unsupervised clustering revealed 13 distinct clusters covering all major cell types in the developing forelimb field. (**E**) Unique gene expression profiles were detected for each major cluster, (**F**) and were largely specific to each cell population.

Unsupervised clustering of the chicken E1.5, E2.5 and E3.5 presumptive forelimb cell populations revealed 13 transcriptionally distinct cell clusters (c) which represented embryonic vasculature and tissues derived from the ectoderm, mesoderm, and endoderm germ layers, covering all major cell types within the presumptive forelimb field (Figure 1c-d). Tissue and cell type identities were assigned to each cluster based on their differential gene expression profiles (observed as differences in log-fold change between clusters; Figure 1e-f) and corresponding spatiotemporal expression patterns observed throughout the developing chicken embryo on the GEISHA chicken gene expression database (Bell et al., 2004; Darnell et al., 2007). Cluster-specific gene expression is shown in Figure 1e-f and Table S1. The embryonic ectoderm comprised two clusters (c2 and c11) defined by expression of *FABP3* and *WNT6*. However, c11 showed unique expression of *FGF8* revealing these cells as progenitors that contribute to formation of the Apical Ectodermal Ridge (AER). The embryonic endoderm (c7) was defined by unique expression of *GUCA2B* and *TTR,* as well as *SHH* (not observed in limb bud mesenchyme due to the early stages sampled). Embryonic vasculature (c8) showed unique expression of *CDH5,* and red blood cells (c13) expressed haemoglobin subunit *HBBA*.

The embryonic mesoderm was found to contribute the largest overall number of cells and was defined by robust expression of *PRRX1*. The mesoderm was comprised of six clusters (c1, c3, c4, c5, c10 and c12) which represented known mesodermal tissue-types during development, which were separated by embryonic stage (Figure 1b, c). Namely, the primitive mesoderm / LPM (c4) was comprised of E1.5 cells and displayed strong expression of early markers such as *MSGN1, EVX1* and *CDX4*. Cells of the E2.5 mesoderm were comprised of clusters representing the somatic LPM (c3) which displayed high *PRRX1* and low *TBX5* expression, splanchnic LPM (c5) with *COLEC10* expression, and the paraxial mesoderm (c1) which displayed unique expression of *TCF15* (Figure 1d, e). Finally, E3.5 cell clusters represented more differentiated tissues, such as the extraembryonic LPM / amnion (c12) through expression of *AQP1,* and limb bud mesenchyme (c10) through high *TBX5* and *FGF10*. We also detected two clusters (c6 and c9) which possessed ubiquitous expression of ectodermal and mesodermal markers, but also high ribosomal and mitochondrial counts, so defined these cells as low diversity cells and were excluded from subsequent analyses.

### Global receptor ligand signalling throughout the differentiating mesoderm

Specification of the mesoderm during embryogenesis is influenced by dynamic intrinsic and extrinsic signalling between the surrounding germ layers and developing tissues. Particularly, members of the BMP, FGF, HH and WNT signalling pathways are known to influence communication between the germ layers (Loh et al., 2016), though the precise ligands and receptors that facilitate different aspects of mesoderm development are unclear. We therefore examined global signalling patterns and ligand-receptor crosstalk between the global cell types during specification of the forelimb field from the aLPM using CellChat (Jin et al., 2021). This analysis revealed extensive signalling pathway usage between different cell types and tissues (Figure 2a), which was altered during developmental progression of key tissue types. The E1.5 primitive MES / LPM (c4) showed the highest level of signalling pathway activity among the tissues studied, with active signalling through more than half of the predicted pathways. This included signalling through the important non-canonical WNT, FGF, HH and BMP pathways, as well as Midkine (MK) and pleiotrophin (PTN), EphrinB, Semaphorin (SEMA3,6), chemokine ligand (CXCL) and adhesion factors laminin, JAM and NECTIN. However, pathway usage was observed to significantly change during subdivision of the aLPM into somatic and splanchnic LPM by E2.5, including decreases in ncWNT, FGF, CXCL and Semaphorin signalling. Splanchnic LPM (c5) formation featured decreased BMP and enhanced AGRN, FN1, CD99 and Ephrin signalling pathways. In contrast, formation of the somatic LPM (c3) was characterised by activated collagen, maintained BMP and NECTIN signalling, and decreased HH signalling. Finally, development of the limb bud mesenchyme (c10) by E3.5 saw activation of WNT and FGF signalling, confirming known interactions, as well as strong activation of Ephrin A and ANGPTL. These observations indicate that development of the LPM is highly dynamic, involving diverse signalling crosstalk during specification and subsequent differentiation (Figure 2a).

**Figure 2.**
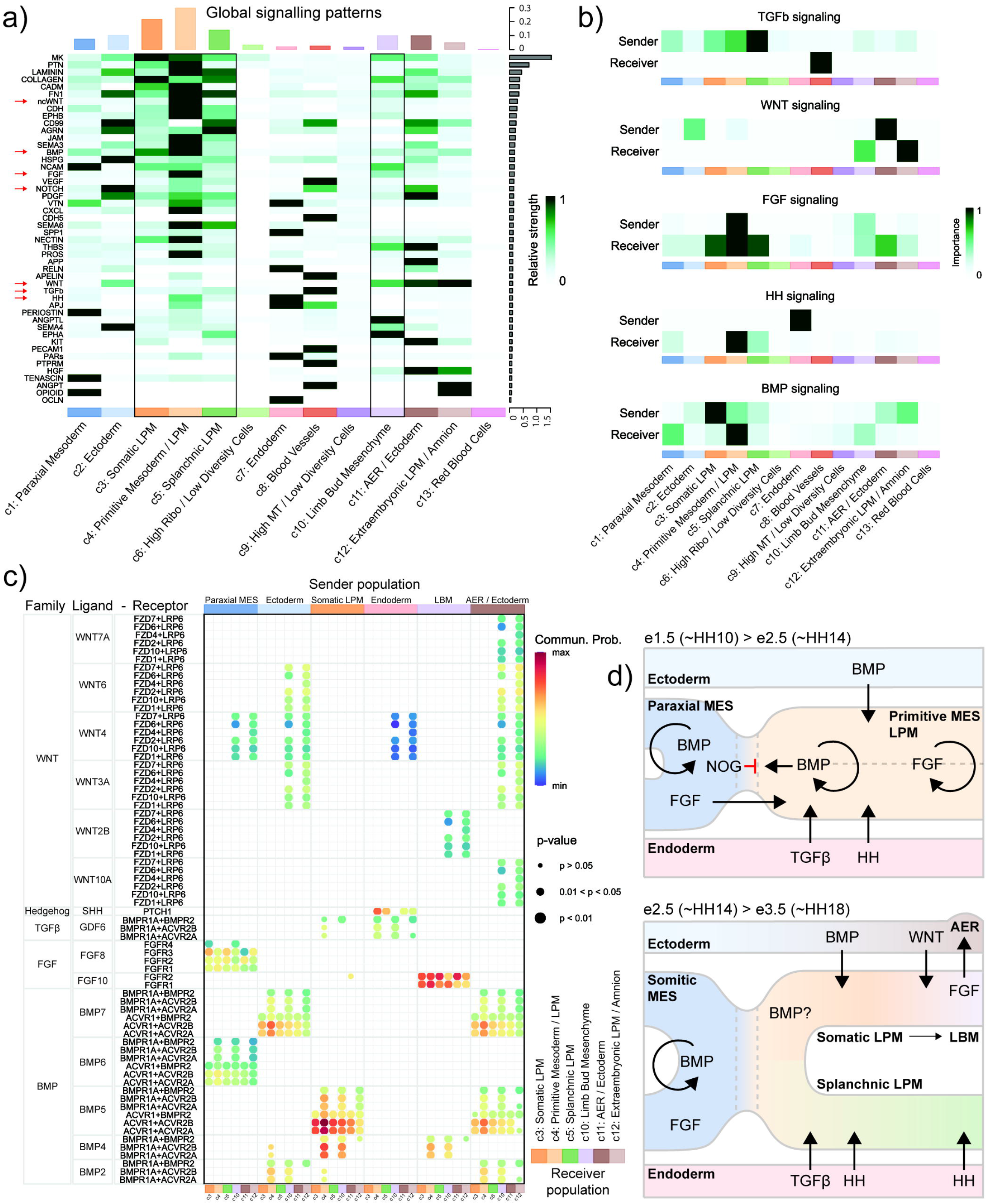
Cellular signalling and ligand-receptor crosstalk in the forelimb field. (**A**) Predictions of active signalling pathways utilized by cell type clusters revealed diverse pathway usage and enriched signalling in the early mesoderm, LPM, ectoderm and limb bud. Major signalling pathways are highlighted by red arrows, emphasizing tissue-specific differences in pathway usage. (**B**) Identification of sender and receiver cell types utilizing major signalling pathways. TGFβ signalling was enriched between the splanchnic LPM and vasculature, WNT signalling between the limb and ectoderm, FGF in the early mesoderm, HH in the endoderm and BMPs in the LPM and ectoderm. (**C**) Identification of key ligand-receptor pairs facilitating tissue-specific signalling from major signalling pathways. This revealed broad, tissue-specific patterns of ligand and receptor heterodimer usage between cell types. For example, BMP7 was identified as the main ligand facilitating ectoderm-LPM signalling, while several WNTs were expressed between the ectoderm and limb bud. FGF10 was confirmed to signal between the limb and ectoderm. (**D**) Diagrammatic summary of signalling pathways active between tissues in the developing limb field.

LPM development and subdivision are thought to be driven by extrinsic ectoderm and endoderm signalling (Roberts et al., 1995; Funayama et al., 1999). We therefore interrogated the datasets to identify the sender and receiver cell-types enriched for TGF-β, WNT, BMP, FGF and HH signalling pathways. This allowed the construction of early signalling networks (Figure 2b). TGF-β did not appear to play a role during LPM differentiation, only received by the embryonic vasculature. The ectoderm was identified as a subtle sender of BMP signals, predicted to be received by the early LPM (c4) and somatic LPM / limb bud (c10), and strong sender of WNT signals which were predicted to be received by the E3.5 limb bud mesenchyme (c10) and amnion (c12). The somatic LPM was also predicted as a strong sender and subtle receiver of BMP signals, though whether this signalling is via an autocrine or paracrine signalling response is unknown. The early mesoderm / LPM was identified as a strong sender and receiver of FGF signals, suggesting intrinsic signalling. The limb bud mesenchyme was additionally identified as a sender of FGF signalling, predicted to be received by the AER ectoderm, confirming the known role of secreted FGF signalling during limb development (Ohuchi et al., 1997). Importantly, the AER ectoderm (c11) was not identified as a sender of FGF signals, despite its important role in establishing the FGF10-FGF8 feedback loop. We predict that this is an artefact of early sampling prior to its activation, as we did not observe enriched *FGF8* expression in the AER cluster (Table S1). Finally, the endoderm was identified as a significant sender of HH signalling, predicted to be received by early LPM, splanchnic LPM and paraxial mesoderm, reaffirming known interactions during splanchnic LPM development and gut formation (Roberts et al., 1995).

With key signalling pathway events established between sender and receiver populations during LPM differentiation and limb development, we next examined which BMP, FGF, HH and WNT ligand and receptor pairs were facilitating these dynamic events (Figure 2b). CellChat (Jin et al., 2021) was further utilized to predict the ligand-receptor crosstalk for each sender-receiver pair of interest, revealing disparate patterns of ligand and receptor usage amongst the different cell populations. *FGF8* was significantly enriched during primitive LPM differentiation, through activation of *FGFR1-4*. During LPM subdivision, endoderm-mesoderm HH signalling occurred exclusively through *SHH* activation of *PTCH1*, with potential contributions by *GDF6* and *BMPR*-*ACVR* receptors in the primitive mesoderm. Ectoderm-mesoderm BMP signalling was seen to be achieved through secretion of BMP7 with activation of combinations of *ACVR1* and *BMPR2*, *ACVR2A* or *ACVR2B* receptor heterodimers. Interestingly, *BMP2* and *BMPR1A* interactions appeared to be only active in the early mesoderm, despite suggested to play a role in LPM subdivision (Funayama et al., 1999). Interestingly, while our data confirm an active role of *BMP4* in early LPM differentiation (Tonegawa et al., 1997) we also observe a significant, yet undefined role of localized *BMP5* signalling during LPM specification, subdivision, and development of the limb bud mesenchyme. Conversely, we see broad expression of WNT ligands in tissues throughout the limb field, including ectodermal *WNT3A, 4, 6, 7A, 10A*, but restricted expression of *FRZD* and *LRP6* receptors only within limb bud mesenchyme and amnion (extraembryonic LPM). Together, these data reveal that signalling throughout the limb field is achieved through combinations of both ubiquitous and tissue-specific ligand-receptor expression patterns (summarized in Figure 2d). Molecular signalling during limb development appears to be largely driven through ectodermal BMP and WNTs, and localized FGFs. Importantly though, the transcriptional targets of these signalling pathways remain largely unknown.

### Specification and differentiation of the LPM

Differentiation of the mesoderm into mature organ and tissue types involves hierarchies of transcriptional regulation (Prummel et al., 2020). However, the early genetic regulators which orchestrate cell fate choices during early mesoderm and LPM differentiation are less clear. We next examined lineage specification and transcriptional dynamics within the LPM by sub-setting the dataset to cells of mesodermal origin. Mesodermal cells were reprocessed with UMAP dimension reduction and Leiden clustering, allocating the 6 previous clusters (c1, c3, c4, c5, c10 and c12; Figure 1a) into 12 new mesoderm sub-clusters (mc1-12; Figure 3a). These captured the previously determined cell types, as well additional intermediate, transitional, and terminal cell types which were altered during progression of developmental stage (Figure 3b). Namely, these updated clusters labelled cells from the somitic (mc9) and intermediate mesoderm (mc10), primitive mesoderm (mc2), early (undifferentiated) LPM (mc1 and mc7), somatic (mc3) and splanchnic (mc4) LPM, mesodermal visceral precursors (mc11), limb bud mesenchyme (mc5), non-limb flank LPM (mc12), extra-embryonic LPM (mc8) and amnion (mc6; Figure 3a). Diagnostic cluster gene markers and log-fold changes are listed in Table S2.

**Figure 3.**
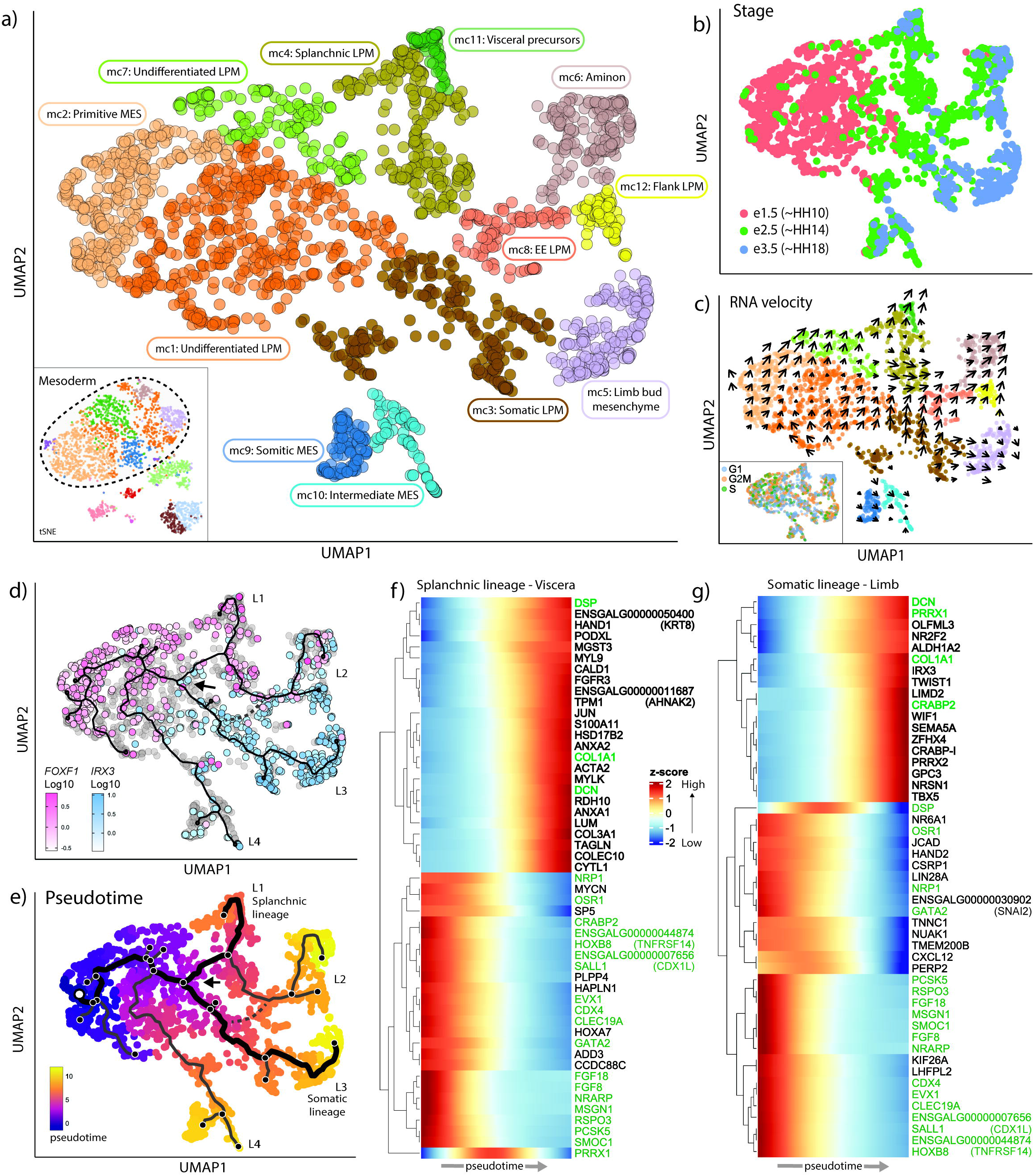
Lineage reconstruction and gene expression dynamics underlying LPM differentiation. (**A**) Subsampling, UMAP projection and re-clustering of mesodermal cell types revealed 12 distinct sub-clusters (sc1-12) with greater resolution of lineage choices and transitional cell types in the forelimb mesoderm, which clearly separated by stage (**B**). (**C**) Transcriptional trajectories were identified through estimates of RNA velocity, revealing distinct lineages of differentiation across different stages of the cell cycle (insert). (**D**) Trajectory inference was computed with Monocle3 further describing 4 lineages (L1, L2, L3, L4) of differentiation. The somatic-splanchnic LPM bifurcation (black arrow) was calibrated using expression of known markers *IRX3* and *FOXF1*, respectively. (**E**) The root node was set in the primitive MES (white circle), and pseudotime calculated to identify gene expression dynamics along the splanchnic and somatic LPM lineages (bold lines). Gene expression dynamics were calculated along the splanchnic (**F**) and somatic (**G**) LPM lineages, revealing activation and repression of distinct gene modules accompanying their differentiation pathways. Unique lineage-specific genes in shown in black while shared genes are in green.

To improve identification of tissue differentiation pathways throughout the LPM, transcriptional dynamics between neighbouring cells of the mesoderm were explored using estimates of RNA velocity. Cell velocity estimates revealed specific, directional transcriptional trajectories between cells as they transitioned from undifferentiated E1.5 (~HH10) precursors towards their distinct tissue fates, connecting three of the four E3.5 (~HH18) cell populations (Figure 3b, c). Interestingly, we identified a heterogeneous population of undifferentiated LPM cells with low directional velocity compared with other neighbouring clusters, despite existing in various stages of the cell cycle (Figure 3c). This suggests that cells of the early LPM may exist in a transiently uncommitted state, before rapidly committing towards a somatic and splanchnic LPM fate by E2.5 (~HH14), likely in response to changes in secreted ectodermal/endodermal signals (Figure 2d). Furthermore, the somitic and intermediate cell clusters of the paraxial mesoderm, despite forming from the primitive mesoderm, did not show a continuum of directional RNA velocities from these precursors. As we did not intentionally sample paraxial tissues, it is unclear whether there were not enough cells isolated to represent the complete differentiation trajectory, or whether these cells possess an earlier embryonic origin to the primitive mesoderm cells captured in our data. As such, we chose to not include this lineage in subsequent analyses.

With transcriptional velocities established, we looked to further define the pathways of differentiation that arise throughout the LPM. The E1.5 primitive mesoderm (mc2) represented the earliest identified cell type so was determined as the root node of mesoderm differentiation. A principal neighbour graph was fit with monocle3 (Trapnell et al., 2014), revealing 4 major lineages (L1-4; Figure 3d) which supported RNA velocity estimates (Figure 3c). These lineages describe the transition from E1.5 (~HH10) primitive mesoderm cells to E3.5 (~HH18) viscera precursors (mc11; L1), non-limb LPM (mc6 and mc12; L2), limb bud mesenchyme (mc5; L3) and somitic/intermediate mesoderm (mc9-10; L4). Importantly, a distinct bifurcation point was observed within the undifferentiated LPM (mc1/mc7), marking LPM subdivision and lineage specification into the somatic (mc3) and splanchnic (mc4) LPM tissue layers (Figure 3d). Subdivision of the undifferentiated LPM is accompanied by localized expression of *IRX3* and *FOXF1* in the somatic and splanchnic LPM, respectively (Funayama et al., 1999; Mahlapuu et al., 2001). To confirm the accuracy of our lineage bifurcation point we examined the expression of *IRX3* and *FOXF1* in the mesoderm. Indeed, cells displayed mutually exclusive expression of *IRX3* or *FOXF1* in complementary domains following the LPM bifurcation point (Figure 3d), with *FOXF1*+ cells observed in the splanchnic mesoderm and viscera precursors (mc4/7/11), and *IRX3*+ cells in the somatopleure, flank LPM and limb bud mesenchyme (mc1/3/5) (Figure 3d), confirming specification of the somatic and splanchnic LPM lineages. However, also included within this lineage graph was an irregular branch linking the splanchnic LPM to the extraembryonic LPM and amnion (Figure 3d, e, dashed line), which does not accurately represent its origins *in vivo*. We were able to resolve this branch through additional k-mean principal graph topology calculations, however these each came at the expense of the somatic-splanchnic LPM bifurcation point, with neither correct topology being present in a single graph without losing cluster resolution (Figure S1). As such, we focused solely on the somatic-splanchnic LPM branching, as confirmed by *FOXF1* and *IRX3* expression, for subsequent lineage trajectory analysis.

### Gene expression dynamics underlying aLPM specification and limb development

Using our reconstructed LPM differentiation lineages, we examined key gene expression dynamics underlying subdivision of the somatic and splanchnic LPM and specification towards a limb or viscera fate. This was achieved through calculations of pseudotime along the viscera and limb lineages (Figure 3e) (Trapnell et al., 2014), where genes with significant expression changes along each branch were identified through Moran’s I spatial autocorrelation (Table S3,S4). The top differentially regulated genes along a given lineage were visualized through expression heatmaps, which grouped these into gene co-activation or repression modules across pseudotime. This revealed a pseudo-temporal hierarchy of dynamically expressed genes during differentiation from the early mesoderm towards a visceral (splanchnic LPM; Figure 3f, Figure S2) or limb (somatic LPM; Figure 3g, Figure S3) fate. Each lineage shared co-expression of primitive mesoderm and early LPM gene modules preceding subdivision, followed by activation of shared or lineage-specific genes and gene modules (Figure 3f, g). During splanchnic LPM differentiation and mesodermal viscera development, ~9 modules of genes were co-activated, including proximal activation of transcription factors *GATA6, NKX2-3, TCF21* and *HAND1*, and repression of *PRRX1* (Figure 3f, Figure S3). Somatic LPM differentiation featured co-activation of ~6 transcriptionally distinct modules, including maintained *PRRX1*, proximal activation of *OLFML3*, *NR2F2* and retinoic acid synthesis *ALDH1A2* (Raldh2), followed by transcription factors *IRX3, IRX6* and bHLH factor *TWIST1* (Figure 3g, Figure S3). Proceeding this was the onset of limb initiation characterised by activation of the limb initiation factor *TBX5,* as well as *TBX2* and *IRX6*, followed by a module of early limb bud mesenchyme genes *LMX1B, PRRX2, WIF1* and *FGF10*. Interestingly, this analysis revealed rapid activation of *TWIST1* prior to limb initiation, suggesting an uncharacterized role during somatic LPM differentiation and early limb initiation.

### TWIST1 in somatic LPM initiation and EMT

Limb initiation has been proposed to occur through a localized EMT of the somatic LPM, dependent on *TBX5* and *FGF10* (Gros and Tabin, 2014). However, prior to the activation of *TBX5* and *FGF10* in the somatic LPM, we observe activation of *PRRX1* and *TWIST1* (Figure 3g) which are known EMT inducers during development and cancer (Ocaña et al., 2012; Fazilaty et al., 2019). We therefore investigated whether these transcription factors may play a role during EMT and initiation of the chicken forelimb. Combined pseudotime gene expression and *in situ* hybridization confirmed *PRRX1* was present in the aLPM forelimb field as early as E1.5/HH10, before becoming regionalized and strongly expressed in the somatic LPM and limb bud (Figure 4a). In comparison, *TWIST1* was detected in the somatic LPM at ~E2.0/HH12-13, but before *TBX5* which was activated shortly after at ~E2.5/HH14 (Figure 4a).

**Figure 4.**
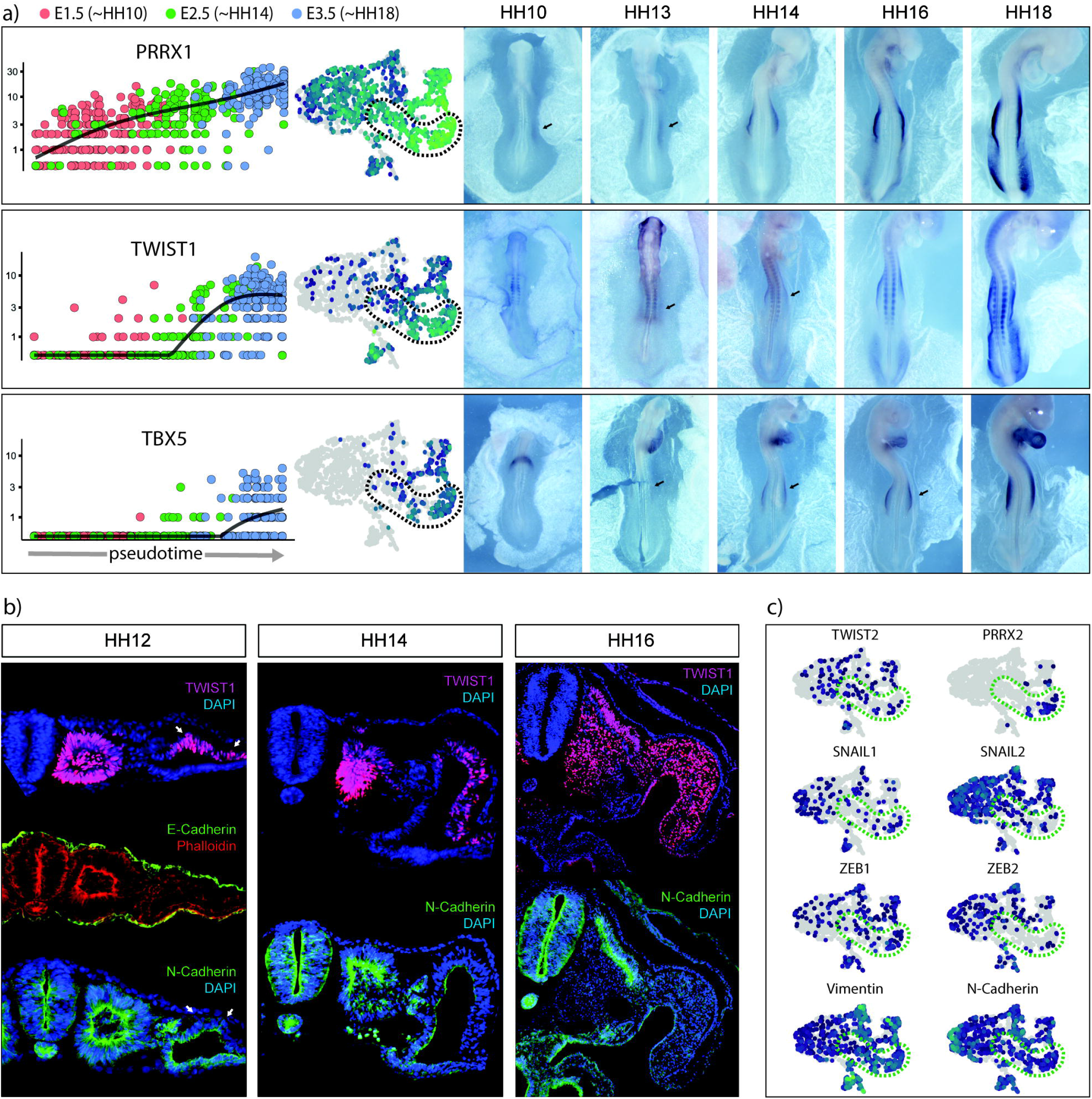
TWIST1 is a likely regulator of somatic LPM and limb bud EMT. (**A**) Expression dynamics of *PRRX1*, *TWIST1* and *TBX5* during somatic LPM lineage specification during pseudotime, and their spatiotemporal *in situ* gene expression profile during early chicken development. *PRRX1* demarks formation and development of the LPM, *TWIST1* is activated in the somatic LPM immediately prior to the onset of *TBX5*. (**B**) Immunofluorescent labelling in the developing forelimb field revealed the LPM possesses mesothelial characteristics, shown by absence of E-Cadherin and presence of N-Cadherin. TWIST1 is observed in the stage (HH) 12 somatic LPM after subdivision, but prior to EMT and proliferation of the limb bud mesenchyme by stage (HH) 16. Note, TWIST1^+^ cells are observed migrating out of the somatic LPM cell layer (arrows). (**C**) *TWIST1* (and *PRRX1*) appears as the major candidate underlying somatic LPM EMT, due to lack of enrichment of other EMT transcription factors in the somatic LPM.

The molecular role of TWIST1 was further examined in the somatic LPM through immunofluorescence. Protein localization in the HH12 forelimb field revealed that the somatic LPM displayed meso-epithelial characteristics expressing N-cadherin, but not E-cadherin (which was restricted to the ectoderm and endoderm) (Figure 4b). Strong TWIST1 expression was detected specifically in the somatic LPM and somites, with some TWIST1+ cells appearing to delaminate from the somatic LPM. By E2.5/HH14 the somatic LPM had undergone proliferation, observed by increased numbers of TWIST1+ cells with reduced N-cadherin, which became localized to the apical edge of the somatic LPM. By ~E3.0/HH16, the forelimb bud was distinct, populated by increased numbers of proliferative TWIST1+ cells (Figure 4b). We also examined whether other EMT transcription factors are expressed during somatic LPM differentiation, but did not detect enrichment of other marker genes, compared to *TWIST1,* in somatic LPM or limb bud mesenchyme clusters (Figure 4c), or along the trajectory (Figure 3c, Figure S3). Rather, the early co-expression of *PRRX1* and *TWIST1* in somatic LPM (Figure 4a, b) suggests that they may play a co-operative role in somatic LPM EMT prior to TBX5-induced limb initiation and outgrowth.

### BMP signalling inhibition disrupts LPM differentiation and limb development

With early markers of somatic LPM defined, we examined how the previously determined signalling interactions influence the expression of early LPM markers. Especially, we looked to define the role of ectodermal BMP signalling during LPM formation, subdivision, and limb bud differentiation, which have been implicated in early somatic LPM cell fate (Funayama et al., 1999). First, we examined whether inhibition of ectodermal-derived BMPs could perturb LPM subdivision and limb initiation. Surgical removal of the forelimb field ectoderm was sufficient to both decrease the developing somatic LPM and reduce *PRRX1* expression (Figure S4a) but was disruptive to development of the embryo and resulted in low viability. Alternatively, we performed targeted electroporation of the secreted BMP antagonist *NOGGIN* into the ~HH9 forelimb field ectoderm to inhibit BMP signalling between the ectoderm and LPM. BMP signal inhibition was not sufficient to disrupt LPM subdivision, despite being previously suggested to drive the process (Funayama et al., 1999). In the absence of BMP signals, the somatic and splanchnic LPM still formed and *PRRX1* and *FOXF1* expression was detected in their respective LPM layers (Figure 5a, Figure S4a). However, inhibition of BMP signalling greatly decreased the overall proportion of somatic LPM cells which formed and had accompanied reduction of *PRRX1* expression compared to the un-electroporated side and GFP controls (Figure 5b). Strikingly, this also significantly inhibited initiation and outgrowth of the developing forelimb bud, seen by significant reductions in *TBX5* and *FGF10* in the somatic LPM and early limb bud (Figure 5a, Figure S4b). Interestingly though, while ectodermal BMP inhibition reduced proportions of somatic LPM cells, it did not appear to influence expression of RA-synthesis gene *ALDH1A2* (Figure S4b) or *TWIST1*, which showed similar expression and protein localization in forelimb sections, albeit in a reduced population of somatic LPM cells compared with controls (Figure 5a, b). Together, these data clarify that ectodermal-mesodermal BMP signalling crosstalk is not necessary for LPM subdivision, but is required for somatic LPM identity, proliferation and early limb initiation and outgrowth. This is achieved through BMP-induced activation of LPM markers including *PRRX1, TBX5* and *FGF10*, but not *ALDH1A2* or *TWIST1* which are some of the first genes activated in the somatic LPM. Thus, while BMP signalling is necessary for somatic LPM development, it appears to be dependent on multiple signalling inputs.

**Figure 5.**
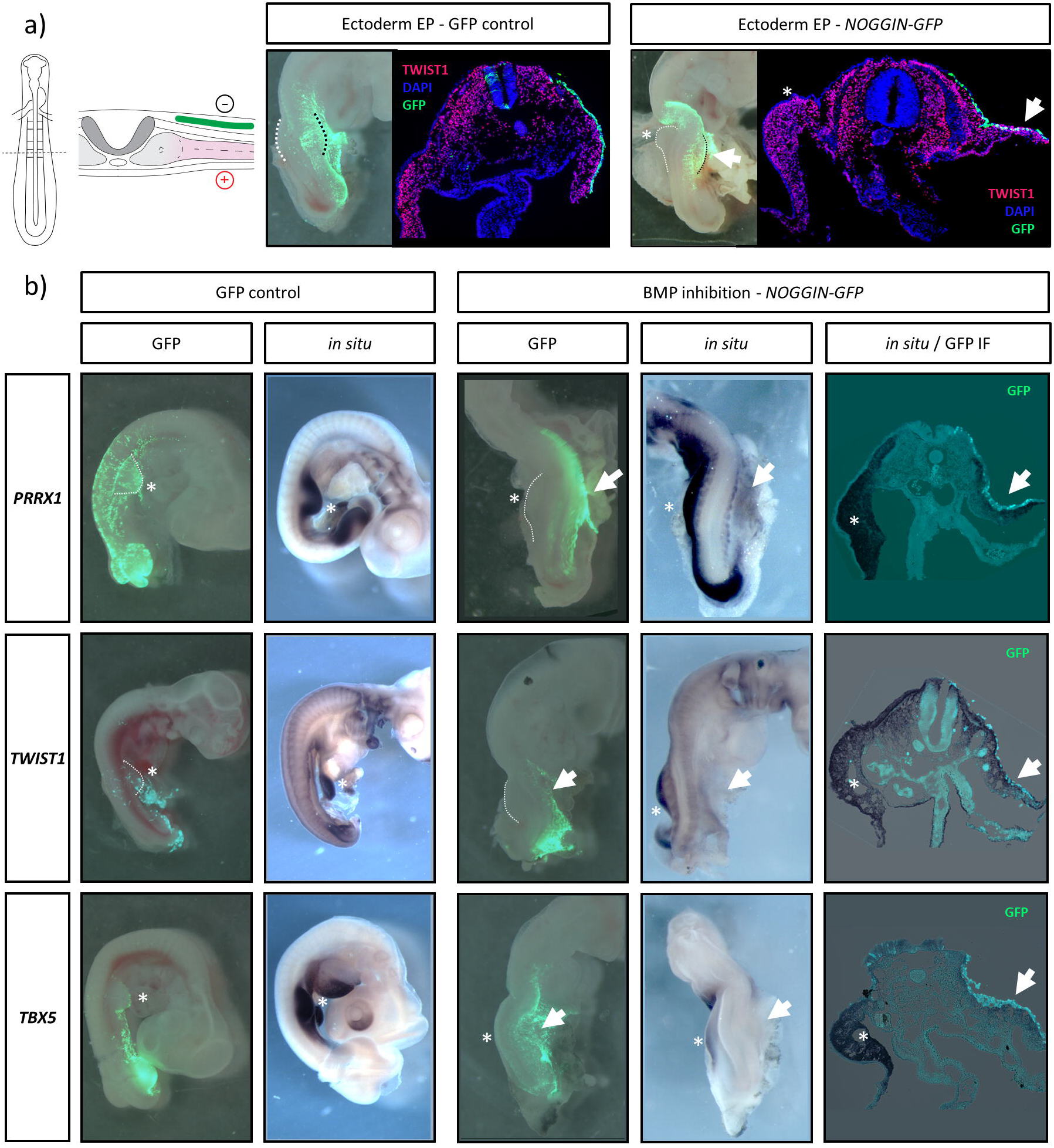
Inhibition of ectodermal BMP signalling severely impacts limb bud development. (**A**) Targeted electroporation of the e1.5 (~HH10) limb field ectoderm. Electroporation of GFP into the ectoderm has no effect on limb bud outgrowth, while BMP antagonism via NOGGIN-GFP greatly inhibits formation of the somatic LPM and possess a greatly reduced limb bud, observed by reduced number of TWIST1+ cells (white arrow). (**B**) Ectodermal-mesodermal BMP antagonism by NOGGIN-GFP decreases activation of *PRRX1* and *TBX5* in the somatic LPM and developing limb bud, as well as in limb bud tissue sections. Note, *TWIST1* does not appear to be as significantly affected by NOGGIN-GFP, as though expression appears reduced in treated limb buds, levels of mRNA expression in limb sections appear unchanged compared to the control side, similar to observations of protein localization (A).

## Discussion

The lateral plate mesoderm gives rise to a wide range of mature tissue types, yet the early developmental events underlying its diversification remain largely undefined. Here, we apply single-cell transcriptomics to investigate cell lineage specification of the early aLPM to define differentiation pathways which direct subdivision of the LPM and specification towards a limb or viscera cell fate. Mesodermal specification originated in a primitive precursor cell type and followed four distinct differentiation pathways from E1.5 (~HH10) to E3.5 (~HH18). Initially, early E1.5 LPM progenitor cells displayed a large degree of cellular heterogeneity (Figure 3a-c) accompanied by extensive signalling pathway usage (Figure 2). This heterogeneity however was rapidly resolved by E2.5, where cells showed commitment to defined differentiation lineages accompanied by dynamic changes in signalling pathway activation and repression. Namely, specification of LPM was achieved through novel ligand-receptor interactions within known BMP, WNT and FGF signalling pathways during somatic LPM and limb formation, and HH signalling in splanchnic LPM development (Roberts et al., 1995; Funayama et al., 1999; Loh et al., 2016), which were distinct by E3.5 (Figure 2). Taken together, the incipient aLPM appears to form in a transient stem-like state, where cells are transcriptionally primed to respond to rapid changes in the extrinsic signalling environment to initiate LPM lineage specification.

LPM subdivision and somatic LPM development are not well defined (Prummel et al., 2019, 2020), but have been suggested to occur through intrinsic activation of *PRRX1* and *IRX3,* and repression of *FOXF1,* in response to ectodermal BMP signalling (Funayama et al., 1999; Mahlapuu et al., 2001; Ocaña et al., 2012). Using pseudotime trajectory analyses, we have defined the temporal activation and repression of gene modules which complement LPM subdivision and differentiation of the somatic and splanchnic LPM lineages (Figure 3f-g, Figure S2,3). We observe activation of several groups of transcription factors which may play critical roles in LPM cell fate. Of note, the earliest stages of LPM subdivision saw repression of primitive mesodermal gene modules and lineage-specific activation of the basic helix-loop-helix (bHLH) transcription factors *HAND1* and *TWIST1* within the splanchnic and somatic LPM lineages, respectively. *HAND1* and *TWIST1* possess known roles in gut and limb development (Firulli et al., 1998; Wu and Howard, 2002; Krawchuk et al., 2010; Loebel et al., 2012), but are yet to be examined during LPM subdivision. Interestingly, bHLH transcription factors form combinations of homo- and heterodimers with unique binding partners to dynamically regulate different biological processes (Fan et al., 2020). We observe expression of both ubiquitous and lineage-specific bHLH binding partners (e.g. *HAND2* and *PRRX1,* respectively) (Fan et al., 2020) throughout the mesodermal cell types. Thus lineage-specific activation of *TWIST1* and *HAND1* may mediate complementary, yet distinct roles in target gene activation during LPM subdivision and differentiation.

The first gene module activated during somatic LPM differentiation included the RA synthesis gene *ALDH1A2* and RA-responsive transcription factor *NR2F2* (Figure 3, S3), suggesting an immediate RA response not captured in our global signalling analysis. Proceeding this was activation of a module including *IRX3* and *TWIST1,* revealing these as the first transcription factors activated along the limb lineage. While *IRX3* is known to be active in the early somatic LPM (Funayama et al., 1999), its role in limb development is not well understood, and deletion of *Irx3* does not produce an overt phenotype in mouse. However, *Irx3/Irx5* double knockout mice have severe limb and heart dysmorphia, suggesting functional co-requirement or redundancy. *TWIST1* has known expression in the somatic LPM (Gitelman, 1997; Tavares et al., 2001), and important roles in limb patterning (Krawchuk et al., 2010; Loebel et al., 2012) though an early role during early limb initiation has not been reported. Limb initiation is posited to begin with EMT of the somatic LPM, dependant on *TBX5* and *FGF10* (Gros and Tabin, 2014). However, *TWIST1* is an important regulator of EMT in development and cancer (Fazilaty et al., 2019) suggesting it may possess an uncharacterized role during early EMT or proliferation of the forelimb field mesenchyme. In support of this idea, specific expression of *TWIST1+* cells in the somatic LPM of HH12 chicken embryos was detected before and after EMT and proliferation of the limb bud mesenchyme (Figure 4). *TWIST1* elicits different biological functions and activity thresholds through dimerization with other transcription factors (Krawchuk et al., 2010; Loebel et al., 2014; Fan et al., 2020), which notably includes the early LPM marker *PRRX1* (Fan et al., 2021). Indeed, this role is supported by observations that *prrx1a* and *twist1b* cooperatively drive EMT and migration of the LPM in zebrafish, which can be rescued by *twist1b* knockdown (Ocaña et al., 2012). Furthermore, *Twist1* null mutant mice possess severely atrophied forelimb buds, potentially through reduced EMT of LPM precursors (Chen and Behringer, 1995). Together, these data strongly implicate *TWIST1* as an early mediator of somatic LPM specification and EMT, though its role in limb initiation is unclear. Intriguingly, an interaction between *TWIST1* and *TBX5* may exist to influence limb initiation but requires further validation.

Our analyses of global ligand-receptor signalling confirmed ectodermal BMP signalling, via BMP2 and BMP7 ligands, as the major active signalling pathway with LPM (Figure 2), supporting its proposed importance in establishing somatic LPM identity (Funayama et al., 1999). To link whether these extrinsic signals were necessary for subdivision of the LPM and activation of intrinsic somatic LPM genes, such as *PRRX1, TWIST1* and *TBX5,* we utilized targeted inhibition of ectodermal-mesodermal BMP signalling via antagonism by NOGGIN. Antagonism of BMP signalling was sufficient to downregulate *PRRX1* in the LPM, but was not necessary for LPM subdivision (Figure 5, S4) as previously suggested (Funayama et al., 1999). However, BMP antagonism produced additional inhibitory effects on limb development through severe atrophy of limb bud outgrowth through a reduced the proportion of somatic LPM cells and complementary reduction of *TBX5* and *FGF10*. Previous studies have revealed that TBX5 activation is achieved through multiple signal inputs, namely *HOX* expression, RA and β*-Catenin/TCF/LEF* through the canonical WNT signalling (Nishimoto et al., 2015). However, while our data shows no evidence for active WNT signalling in the somatic LPM (Figure 2), it establishes a novel role of BMP signalling in *TBX5/FGF10* activation. Notably, inhibition of BMP signalling prior to its critical window of activity between the ectoderm and early LPM is sufficient to inhibit forelimb outgrowth through reduced commitment of somatic LPM precursors. This is in contrast to previous studies utilizing retroviral delivery of *NOGGIN* to somatic LPM, which produce smaller limbs with patterning defects through aberrant formation of the AER (Capdevila and Johnson, 1998; Pizette and Niswander, 1999). Additionally, BMP signalling was not sufficient to influence TWIST1 in the somatic LPM but instead may be RA responsive. RA pathway genes *ALDH1A2* and *NR2F2* were activated in the LPM immediately prior to TWIST1 activation (Figure 3, S3), and *TWIST1* can be induced during limb bud outgrowth through ectopic addition of RA (Tavares et al., 2001). However, the influence of RA in *TWIST1* induction in the early somatic LPM has yet to be seen. Nevertheless, these data suggest a model where LPM subdivision and somatic LPM differentiation may be achieved through the combined action of BMP and RA signalling.

This study establishes the first transcriptional atlas of progenitor, transitional and maturing cell types throughout the early forelimb field and uncovers the global signalling pathways and key transcription factors that are activated within developing tissues. We have begun to shed light on the early cell fate decisions that initiate development of the vertebrate limb, though additional analyses will strengthen the essential factors underlying its development. Particularly, as our data only captures the earliest stages of limb initiation, integrative analysis with later stages of limb patterning (Feregrino et al., 2019; Feregrino and Tschopp, 2021) provides an opportunity to reconstruct the full cellular and developmental events underlying forelimb initiation, patterning and development. Furthermore, while we shed light on the genetic hierarchy that is active during LPM specification, the gene regulatory networks accompanying these are unknown. Applied single cell ATAC-seq and single-cell gene regulatory network reconstruction (Aibar et al., 2017) would further allowing the construction of gene regulatory networks within the developing LPM. Our data highlight a previously unidentified role for *TWIST1* as an early mediator of somatic LPM development, though additional work is required to define its precise role. Together, the application of these additional analysis will yield greater clarification into the processes that drive development of mesodermal precursors into complex structures such as the vertebrate limb.

## Methods

### Egg incubation, tissue collection, single cell sampling

Chicken eggs were collected at embryonic day (e) 1.5 (stage 10), E2.5 (HH14) or E3.5 (HH18), while emu eggs were collected at E3.5 (HH10) e4.5 (HH14) and e5.5 (HH18). Embryos were dissected away from extra-embryonic membranes, rinsed in ice-cold DPBS then the LPM dissected. LPM tissues were digested with 0.05% Trypsin / EDTA and incubated at 37°C for 15 minutes, with mechanical dissociation every 5 minutes until no clumps were visible. Enzymatic activity was stopped with addition of 10% FCS. The dissociated cells were spun at 400g for 5 minutes, then resuspended in 1x EDTA / Propidium Iodide in DMEM (Gibco). Cells were filtered through a 70um Flowmi Cell Strainer (Scienceware), and viable cells were isolated through flow cytometry (Flowcore, Monash University).

Samples were submitted to Micromon Genomics (Monash University) for analysis using the 10X Genomics Chromium Controller and Chromium Single Cell 30 Library & Gel Bead Kit V2, as per the manufacturer’s instructions. Samples were subjected to 10 cycles of PCR for cDNA amplification and 16 cycles for library amplification. Completed libraries were pooled in an equimolar ratio along with 5% PhilX Control Library V3 (Illumina), denatured and diluted to 2.0pM as per the manufacturer’s instructions. The prepared libraries were sequencing using an Illumina Next-Seq500 using Illumina 150c V2 chemistry and V2.5 flow cell, as per the manufacturer’s instructions.

### Bioinformatics Pre-processing

Reads were aligned to the chicken GRCg6a reference using CellRanger (v4.0.0, using option: –force-cells 15000). Due to the number of reads observed just downstream of annotated genes, the gene annotation (from ensembl release 100, gene biotypes: protein coding, lincRNA and antisense) was edited to include 1000bp downstream each gene. Single cell analysis was performed in R using packages scran (Lun et al., 2021), scater (Mccarthy et al., 2017) for QC and iSEE for interactive viewing (Lun et al., 2018). Gene names were used for analysis and, where they mapped to multiple ensembl ids, the ensembl ID with the highest number of counts was kept. Cells with low total umi counts (<2000) were excluded. Cell cycle was annotated with cyclone in the scran package (Lun et al., 2021) using the mouse reference from (Scialdone et al., 2015) mapped to its one-to-one chicken orthologs.

The top 1000 genes with the highest biological variance were identified with modelGeneVar function of scran (Lun et al., 2021), blocked on the sequencing sample, and excluded mitochondrial genes or genes on the Z or W chromosomes to minimise sex effects. PCA was calculated on these, and the first 15 PCs used to generate a global chicken tSNE layout. Clusters were defined with the walktrap method, on a SNN graph (k=10) (Lun et al., 2021), and cluster identities were determined from gene logFC changes and spatial expression profiles in the Gallus Expression *In Situ* Hybridization Analysis (GEISHA) database (Bell et al., 2004; Darnell et al., 2007).

### Bioinformatic analysis

#### Global signalling pathway usage and ligand-receptor crosstalk

Global signalling patterns throughout the chicken forelimb field were examined using the R package CellChat (Jin et al., 2021), where signalling communication networks were constructed based on 1:1 gene orthology with a curated *Homo sapiens* database and default parameters. Visualizations of pathway and ligand receptor signalling were generated with CellChat and edited with Adobe Illustrator.

#### Mesoderm analysis and lineage reconstruction

Mesodermal cell clusters were subset from the full tSNE for additional, focused analyses. Briefly, mesodermal clusters were subset to a new object, PCA and UMAP dimension reduction was recalculated using the previously determined highly variable genes and corrected for cell cycle effects. Next, the object was imported into Monocle3 (Spielmann et al.; Trapnell et al., 2014) for clustering (k=4). Cluster labels were confirmed by identifying differentially expressed marker genes through regression analysis implemented in monocle *fit_models* function, producing distinct 12 clusters covering all known cell types within the developing limb field mesoderm.

Estimations of RNA velocity were produced, where reads were aligned to the reference a second time with STAR solo (v2.7.5) to identify proportions of spliced and unspliced transcripts (Dobin et al., 2013). Velocity analysis with performed with velocyto.R (La Manno et al., 2018), and directional transcriptional velocities between cells were visualized. Lineage trajectories throughout the mesoderm were additionally constructed in Monocle3 using reverse graph embedding (k=4, minimum branch length = 15, rann.k = 50), which produced 4 major lineages that originated in E1.5 cells and terminated in E3.5 cells. Lineage bifurcation points were corroborated using know LPM marker gene expression through the *plot_cells* function, then pseudotime was calculated by selecting the origin of the lineages using the *order_cells* function. Then, to identify genes that dynamically changed in expression across pseudotime, key lineages throughout the mesoderm were subset using the *chose_graph_segments* function and graph tests were run to identify lineage-specific, differentially regulated genes and filtered based on Moran’s I statistic and q value. Modules of genes that significantly changed across pseudotime were visualized by hierarchical clustering through the R package ComplexHeatmap. Gene expression in individual cells across pseudotime were further visualized using the *plot_gene_in_pseuodtime* function in Monocle3.

### Functional experimentation

#### Gene expression analysis by in situ hybridization and immunofluorescence

Whole mount *in situ* hybridization for spatial mRNA expression was carried out as described previously (Smith et al., 2016) with minor modifications. Briefly, whole HH8-HH18 chicken embryos were fixed overnight in 4% paraformaldehyde, dehydrated in methanol, and rehydrated in PBTX (PBS + 0.1% Triton X-1000). Tissues were permeabilized in 10mg/mL proteinase K for up to 1 hour, depending upon size then re-fixed in glutaraldehyde/ 4% PFA. Tissues underwent pre-hybridization (50% formamide, 5 x SSC, 0.1% Triton X-100, 0.5% CHAPS, 5mM EDTA, 50mg/mL Heparin, 1mg/mL yeast RNA, 2% blocking powder) overnight at 65°C. Riboprobe templates were provided as gifts, generated from public sources, or designed and synthesised in house. Primer sequences and/or source are listed in Table S5. Where applicable, templates were amplified from limb and whole embryo cDNA using gene specific primers. Fragments were resolved by 1% agarose electrophoresis, excised, and purified using a Nucleospin PCR clean-up kit and subcloned into p-GEM T-easy (Promega). Antisense RNA probes were synthesized using T3, T7 or SP6 RNA polymerases and the DIG-labelling kit (Roche, #11277073910) as per the manufacturer’s instructions. Precipitated probes were added to pre-hybridized tissues (approx. 5mL/ tube) and hybridization was carried out overnight at 65°C. Tissues were then subjected to stringency washes, blocked in BSA, then treated overnight with anti-DIG antibody conjugated with alkaline phosphatase. Tissues were exposed to BCIP/NBT colour reaction at room temperature for up to 3 hours (340mg/mL NBT and 175 mg/mL BCIP in NTMT (100mM NaCl, 100mM Tris-HCl, pH9.5, 50mM MgCl2, 0.1% Tween-20).

Chicken embryos were fixed in 4% PFA/PBS for 15 minutes at room temperature then cryo-protected in 30% sucrose. Embryos were snap frozen in OCT and 10mm frozen sections were cut. Antigen retrieval was performed for detection of TWIST1, otherwise sections were left in PBS. For co-detection of TWIST1 and other markers in the LPM, antibody incubations were performed on successive tissue sections. Sections were blocked and permeabilised in 1% Triton X-100, 2% BSA/PBS for 1-2hr at room temperature, then incubated with primary antibody in 0.5% Triton X-100, 1% BSA/PBS incubation overnight at 4°C.

#### Targeted electroporation

Electroporation of chicken ectoderm was performed using custom parameters. Briefly, eggs were incubated for ~36 hours until stage HH8-HH10. Here, a solution containing TOL2 Transposase and NOGGIN-GFP plasmids at final concentrations of 1ug/ul were mixed with 0.1% fast green and injected between the vitelline membrane and embryo. Electroporation was performed by placing the positive electrode above the presumptive forelimb field on the right side of the embryo, and negative electrode under the embryo above the yolk, and delivered through 3x 10V, 60ms width, 50ms space pulses (Intracel TSS20 Ovodyne Electroporator). Eggs were sealed with tape and the embryos were incubated for a further 24 – 48h. Embryos were then harvested and GFP imaged on a Fluorescence dissecting microscope. GFP positive embryos were then fixed in 4% PFA overnight at 4°C.

## Acknowledgements

We thank Micromon Genomics for assistance with sequencing design, Monash University Flowcore for flow cytometry, and Monash Histology platform for histological processing.

We additionally thank Alex Combes and Kieran Short for helpful suggestions for choice of bioinformatics methods, and constructive comments from all members of the Smith Lab. This work was supported by the Australian Research Council Discovery Project scheme (DP190100890 to Craig Smith).

## Author contributions

A.H.N and C.A.S conceptualized and designed the study. A.H.N performed the experiments. A.T.M assisted with flow cytometry. S.M.W performed bioinformatic pre-processing. A.H.N and S.M.W performed computational analysis. A.H.N, A.T.M and C.A.S analysed and interpreted the data. All authors contributed to preparation of the manuscript.

## Declaration of interests

The authors declare no competing interests.

## Notes

### Competing Interest Statement

The authors have declared no competing interest.

## References

Agarwal, P., Wylie, J.N., Galceran, J., Arkhitko, O., Li, C., Deng, C., Grosschedl, R., and Bruneau, B.G. 2003. Tbx5 is essential for forelimb bud initiation following patterning of the limb field in the mouse embryo. Development 130: 623–633.

Aibar, S., González-Blas, C.B., Moerman, T., Huynh-Thu, V.A., Imrichova, H., Hulselmans, G., Rambow, F., Marine, J.C., Geurts, P., Aerts, J., et al. 2017. SCENIC: Single-cell regulatory network inference and clustering. Nat. Methods 14: 1083–1086.

Bell, G.W., Yatskievych, T.A., and Antin, P.B. 2004. GEISHA, a whole-mount in situ hybridization gene expression screen in chicken embryos. Dev. Dyn. 229: 677–687.

Capdevila, J., and Johnson, R.L. 1998. Endogenous and ectopic expression of noggin suggests a conserved mechanism for regulation of BMP function during limb and somite patterning. Dev. Biol. 197: 205–217.

Chen, Z.F., and Behringer, R.R. 1995. Twist Is Required in Head Mesenchyme for Cranial Neural Tube Morphogenesis. Genes Dev. 9: 686–699.

Darnell, D.K., Kaur, S., Stanislaw, S., Davey, S., Konieczka, J.H., Yatskievych, T.A., and Antin, P.B. 2007. GEISHA: an in situ hybridization gene expression resource for the chicken embryo. Cytogenet. Genome Res. 117: 30–35.

Dobin, A., Davis, C.A., Schlesinger, F., Drenkow, J., Zaleski, C., Jha, S., Batut, P., Chaisson, M., and Gingeras, T.R. 2013. Sequence analysis. 29: 15–21.

Fan, X., Pragathi Masamsetti, V., Sun, J.Q.J., Engholm-Keller, K., Osteil, P., Studdert, J., Graham, M.E., Fossat, N., and Tam, P.P.L. 2021. Twist1 and chromatin regulatory proteins interact to guide neural crest cell differentiation. Elife 10: 1–71.

Fan, X., Waardenberg, A.J., Demuth, M., Osteil, P., Sun, J.Q.J., Loebel, D.A.F., Graham, M., Tam, P.P.L., and Fossat, N. 2020. TWIST1 Homodimers and Heterodimers Orchestrate Lineage-Specific Differentiation. Mol. Cell. Biol. 40: 1–20.

Fazilaty, H., Rago, L., Kass Youssef, K., Ocaña, O.H., Garcia-Asencio, F., Arcas, A., Galceran, J., and Nieto, M.A. 2019. A gene regulatory network to control EMT programs in development and disease. Nat. Commun. 10:.

Feregrino, C., Sacher, F., Parnas, O., and Tschopp, P. 2019. A single-cell transcriptomic atlas of the developing chicken limb. BMC Genomics 20: 1–15.

Feregrino, C., and Tschopp, P. 2021. Assessing evolutionary and developmental transcriptome dynamics in homologous cell types. Dev. Dyn. 1–18.

Firulli, A.B., McFadden, D.G., Lin, Q., Srivastava, D., and Olson, E.N. 1998. Heart and extra-embryonic mesodermal defects in mouse embryos lacking the bHLH transcription factor Hand1. Nat. Genet. 18: 266–270.

Funayama, N., Sato, Y., Matsumoto, K., Ogura, T., and Takahashi, Y. 1999. Coelom formation: Binary decision of the lateral plate mesoderm is controlled by the ectoderm. Development 126: 4129–4138.

Gerber, T., Murawala, P., Knapp, D., Masselink, W., Schuez, M., Hermann, S., Gac-Santel, M., Nowoshilow, S., Kageyama, J., Khattak, S., et al. 2018. Single-cell analysis uncovers convergence of cell identities during axolotl limb regeneration. Science (80-.). 362: eaaq0681.

Gitelman, I. 1997. Twist protein in mouse embryogenesis. Dev. Biol. 189: 205–214.

Gros, J., and Tabin, C.J. 2014. Vertebrate Limb Bud Formation Is Initiated by Localized Epithelial-to-Mesenchymal Transition. Science (80-.). 343: 1253–1256.

Hamburger, V., and Hamilton, H.L. 1951. A series of normal stages in the development of the chick embryo. J. Morphol. 88: 49–92.

Han, L., Chaturvedi, P., Kishimoto, K., Koike, H., Nasr, T., Iwasawa, K., Giesbrecht, K., Witcher, P.C., Eicher, A., Haines, L., et al. 2020. Single cell transcriptomics identifies a signaling network coordinating endoderm and mesoderm diversification during foregut organogenesis. Nat. Commun. 11:.

Harvey, R.P., Lai, D., Elliott, D., Biben, C., Solloway, M., Prall, O., Stennard, F., Schindeler, A., Groves, N., Lavulo, L., et al. 2002. Homeodomain factor Nkx2-5 in heart development and disease. Cold Spring Harb. Symp. Quant. Biol. 67: 107–114.

Jin, S., Guerrero-Juarez, C.F., Zhang, L., Chang, I., Ramos, R., Kuan, C.H., Myung, P., Plikus, M. V., and Nie, Q. 2021. Inference and analysis of cell-cell communication using CellChat. Nat. Commun. 12: 1–20.

Johnson, G.L., Masias, E.J., and Lehoczky, J.A. 2020. Cellular Heterogeneity and Lineage Restriction during Mouse Digit Tip Regeneration at Single-Cell Resolution. Dev. Cell 52: 525–540.e5.

Krawchuk, D., Weiner, S.J., Chen, Y.T., Lu, B.C., Costantini, F., Behringer, R.R., and Laufer, E. 2010. Twist1 activity thresholds define multiple functions in limb development. Dev. Biol. 347: 133–146.

Kuratani, S., Martin, J.F., Wawersik, S., Lilly, B., Eichele, G., and Olson, E.N. 1994. The expression pattern of the chick homeobox gene gMHox suggests a role in patterning of the limbs and face and in compartmentalization of somites. Dev. Biol. 161: 357–369.

Li, D., Sakuma, R., Vakili, N.A., Mo, R., Puviindran, V., Deimling, S., Zhang, X., Hopyan, S., and Hui, C. 2014. Formation of Proximal and Anterior Limb Skeleton Requires Early Function of Irx3 and Irx5 and Is Negatively Regulated by Shh Signaling. Dev. Cell 29: 233–240.

Loebel, D.A.F., Hor, A.C.C., Bildsoe, H., Jones, V., Chen, Y.T., Behringer, R.R., and Tam, P.P.L. 2012. Regionalized Twist1 activity in the forelimb bud drives the morphogenesis of the proximal and preaxial skeleton. Dev. Biol. 362: 132–140.

Loebel, D.A.F., Hor, A.C.C., Bildsoe, H.K., and Tam, P.P.L. 2014. Timed deletion of Twist1 in the limb bud reveals age-specific impacts on autopod and zeugopod patterning. PLoS One 9:.

Logan, M., Simon, H.G., and Tabin, C. 1998. Differential regulation of T-box and homeobox transcription factors suggests roles in controlling chick limb-type identity. Development 125: 2825–2835.

Loh, K.M.M., Chen, A., Koh, P.W.W., Deng, T.Z.Z., Sinha, R., Tsai, J.M.M., Barkal, A.A.A., Shen, K.Y.Y., Jain, R., Morganti, R.M.M., et al. 2016. Mapping the Pairwise Choices Leading from Pluripotency to Human Bone, Heart, and Other Mesoderm Cell Types. Cell 166: 451–467.

Lun, A.T.L., Mccarthy, D.J., and Marioni, J.C. 2021. A step-by-step workflow for low-level analysis of single-cell RNA-seq data with Bioconductor [ version 2◻ peer review◻: 3 approved, 2 approved with reservations].

Lun, A.T.L., Rue-Albrecht, K., Marini, F., and Soneson, C. 2018. iSEE: Interactive SummarizedExperiment Explorer [version 1; referees: 2 approved]. F1000Research 7:.

Mahadevaiah, S.K., Sangrithi, M.N., Hirota, T., and Turner, J.M.A. 2020. A single-cell transcriptome atlas of marsupial embryogenesis and X inactivation. Nature 586: 612–617.

Mahlapuu, M., Ormestad, M., Enerbäck, S., and Carlsson, P. 2001. The forkhead transcription factor Foxf1 is required for differentiation of extra-embryonic and lateral plate mesoderm. Development 128: 155–166.

La Manno, G., Soldatov, R., Zeisel, A., Braun, E., Hochgerner, H., Petukhov, V., Lidschreiber, K., Kastriti, M.E., Lönnerberg, P., Furlan, A., et al. 2018. RNA velocity of single cells. Nature 560: 494–498.

Martin, J.F., Bradley, A., Olson, E.N., and Eric, N.O. 1995. The paired-like homeo box gene MHox is required for early events of skeletogenesis in multiple lineages. Genes Dev. 9: 1237–1249.

Mccarthy, D.J., Campbell, K.R., Lun, A.T.L., and Wills, Q.F. 2017. Gene expression Scater◻: pre-processing , quality control , normalization and visualization of single-cell RNA-seq data in R. 33: 1179–1186.

Moon, A.M., and Capecchi, M.R. 2000. Fgf8 is required for outgrowth and patterning of the limbs. Nat. Genet. 26: 455–459.

Newton, A.H., and Smith, C.A. 2020. Regulation of vertebrate forelimb development and wing reduction in the flightless emu. Dev. Dyn. dvdy.288.

Nishimoto, S., and Logan, M.P.O. 2016. Subdivision of the lateral plate mesoderm and specification of the forelimb and hindlimb forming domains. Semin. Cell Dev. Biol. 49: 102–108.

Nishimoto, S., Wilde, S.M., Wood, S., and Logan, M.P.O. 2015. RA Acts in a Coherent Feed-Forward Mechanism with Tbx5 to Control Limb Bud Induction and Initiation. Cell Rep. 12: 879–891.

Ocaña, O.H., Córcoles, R., Fabra, Á., Moreno-Bueno, G., Acloque, H., Vega, S., Barrallo-Gimeno, A., Cano, A., Nieto, M.A., Co, R., et al. 2012. Metastatic Colonization Requires the Repression of the Epithelial-Mesenchymal Transition Inducer Prrx1. Cancer Cell 22: 709–724.

Ohuchi, H., Nakagawa, T., Yamamoto, A., Araga, A., Ohata, T., Ishimaru, Y., Yoshioka, H., Kuwana, T., Nohno, T., Yamasaki, M., et al. 1997. The mesenchymal factor, FGF10, initiates and maintains the outgrowth of the chick limb bud through interaction with FGF8, an apical ectodermal factor. Development 124: 2235–2244.

Peterson, R.S., Lim, L., Ye, H., Zhou, H., Overdier, D.G., and Costa, R.H. 1997. The winged helix transcriptional activator HFH-8 is expressed in the mesoderm of the primitive streak stage of mouse embryos and its cellular derivatives. Mech. Dev. 69: 53–69.

Pijuan-Sala, B., Griffiths, J.A., Guibentif, C., Hiscock, T.W., Jawaid, W., Calero-Nieto, F.J., Mulas, C., Ibarra-Soria, X., Tyser, R.C.V., Ho, D.L.L., et al. 2019. A single-cell molecular map of mouse gastrulation and early organogenesis. Nature 566: 490–495.

Pizette, S., and Niswander, L. 1999. BMPs negatively regulate structure and function of the limb apical ectodermal ridge. Development 126: 883–894.

Prummel, K.D., Hess, C., Nieuwenhuize, S., Parker, H.J., Rogers, K.W., Kozmikova, I., Racioppi, C., Brombacher, E.C., Czarkwiani, A., Knapp, D., et al. 2019. A conserved regulatory program initiates lateral plate mesoderm emergence across chordates. Nat. Commun. 10: 1–15.

Prummel, K.D., Nieuwenhuize, S., and Mosimann, C. 2020. The lateral plate mesoderm. Dev. 147:.

Rallis, C., Bruneau, B.G., Del Buono, J., Seidman, C.E., Seidman, J.G., Nissim, S., Tabin, C.J., and Logan, M.P.O. 2003. Tbx5 is required for forelimb bud formation and continued outgrowth. Development 130: 2741–2751.

Roberts, D.J., Johnson, R.L., Burke, A.C., Nelson, C.E., Morgan, B.A., and Tabin, C. 1995. Sonic hedgehog is an endodermal signal inducing Bmp-4 and Hox genes during induction and regionalization of the chick hindgut. Development 121: 3163–3174.

Scialdone, A., Natarajan, K.N., Saraiva, L.R., Proserpio, V., Teichmann, S.A., Stegle, O., Marioni, J.C., and Buettner, F. 2015. Computational assignment of cell-cycle stage from single-cell transcriptome data. Methods 85: 54–61.

Scialdone, A., Tanaka, Y., Jawaid, W., Moignard, V., Wilson, N.K., Macaulay, I.C., Marioni, J.C., and Göttgens, B. 2016. Resolving early mesoderm diversification through single-cell expression profiling. Nature 535: 289–293.

Smith, C.A., Farlie, P.G., Davidson, N.M., Roeszler, K.N., Hirst, C., Oshlack, A., and Lambert, D.M. 2016. Limb patterning genes and heterochronic development of the emu wing bud. Evodevo 7: 1–17.

Spielmann, M., Qiu, X., Huang, X., Ibrahim, D.M., Hill, A.J., and Zhang, F. The single-cell transcriptional landscape of mammalian organogenesis. Nature.

Tanaka, M. 2016. Developmental Mechanism of Limb Field Specification along the Anterior–Posterior Axis during Vertebrate Evolution. J. Dev. Biol. 4: 18.

Tavares, A.T., Izpisúa-Belmonte, J.C., and Rodríguez-León, J. 2001. Developmental expression of chick Twist and its regulation during limb patterning. Int. J. Dev. Biol. 45: 707–713.

Tickle, C. 2015. How the embryo makes a limb: Determination, polarity and identity. J. Anat. 227: 418–430.

Tonegawa, A., Funayama, N., Ueno, N., and Takahashi, Y. 1997. Mesodermal subdivision along the mediolateral axis in chicken controlled by different concentrations of BMP-4. Development 124: 1975–1984.

Tonegawa, A., and Takahashi, Y. 1998. Somitogenesis Controlled by Noggin. Dev. Biol. 202: 172–182.

Trapnell, C., Cacchiarelli, D., Grimsby, J., Pokharel, P., Li, S., Morse, M., Lennon, N.J., Livak, K.J., Mikkelsen, T.S., and Rinn, J.L. 2014. The dynamics and regulators of cell fate decisions are revealed by pseudotemporal ordering of single cells. Nat. Biotechnol. 32: 381–386.

Wang, Q., Lan, Y., Cho, E.-S., Maltby, K.M., and Jiang, R. 2005. Odd-skipped related 1 (Odd1) is an essential regulator of heart and urogenital development. Dev. Biol. 288: 582–594.

Wu, X., and Howard, M.J. 2002. Transcripts encoding hand genes are differentially expressed and regulated by BMP4 and GDNF in developing avian gut. Gene Expr. 10: 279–293.

Yoshino, T., Murai, H., and Saito, D. 2016. Hedgehog-BMP signalling establishes dorsoventral patterning in lateral plate mesoderm to trigger gonadogenesis in chicken embryos. Nat. Commun. 7: 1–11.

Zuniga, A.A. 2015. Next generation limb development and evolution: Old questions, new perspectives. Dev. 142: 3810–3820.

