## Supplementary figures and images for "Cell lineage specification during development of the anterior lateral plate mesoderm and forelimb field"

### Figure S1

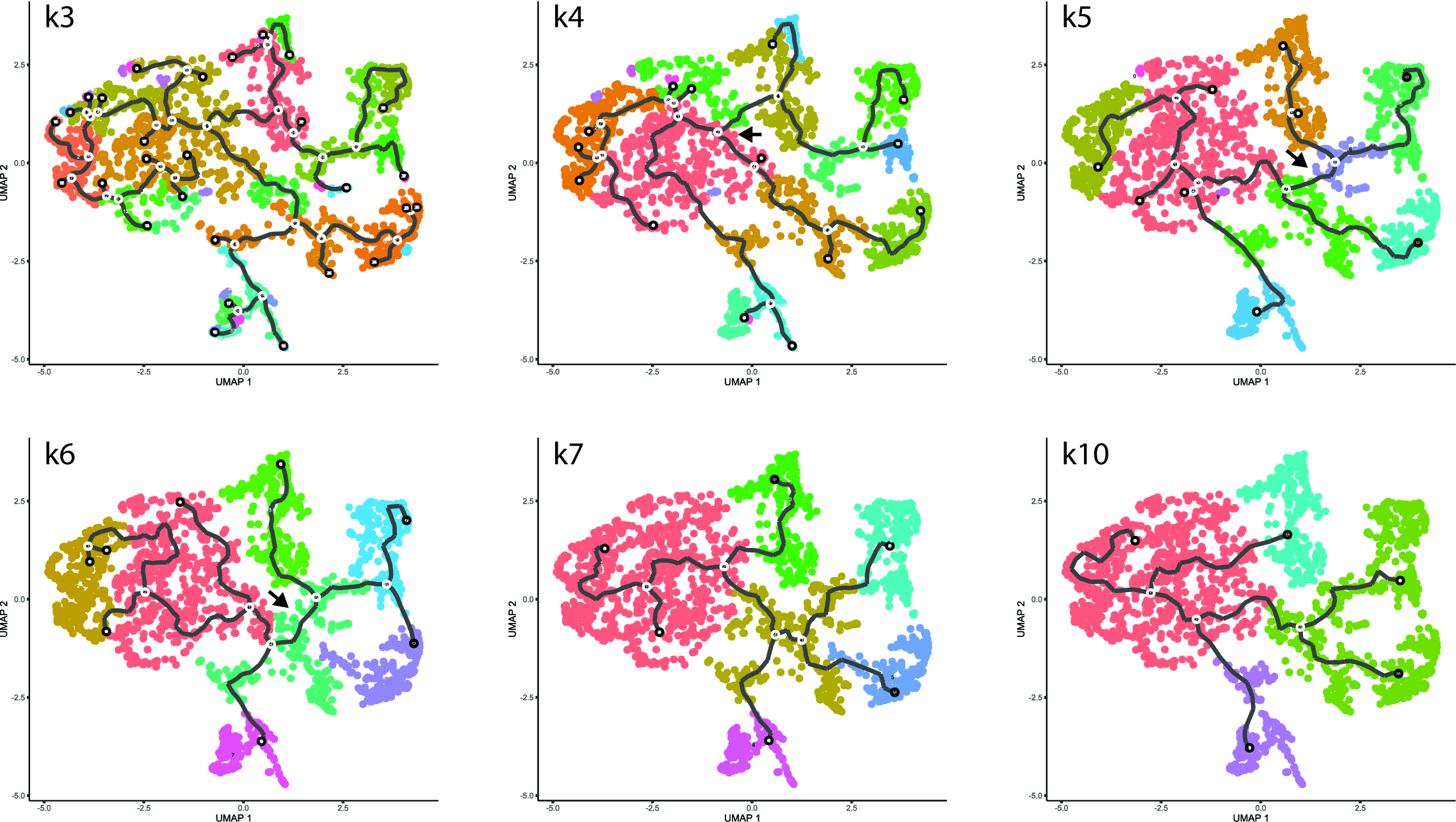

### Figure S2

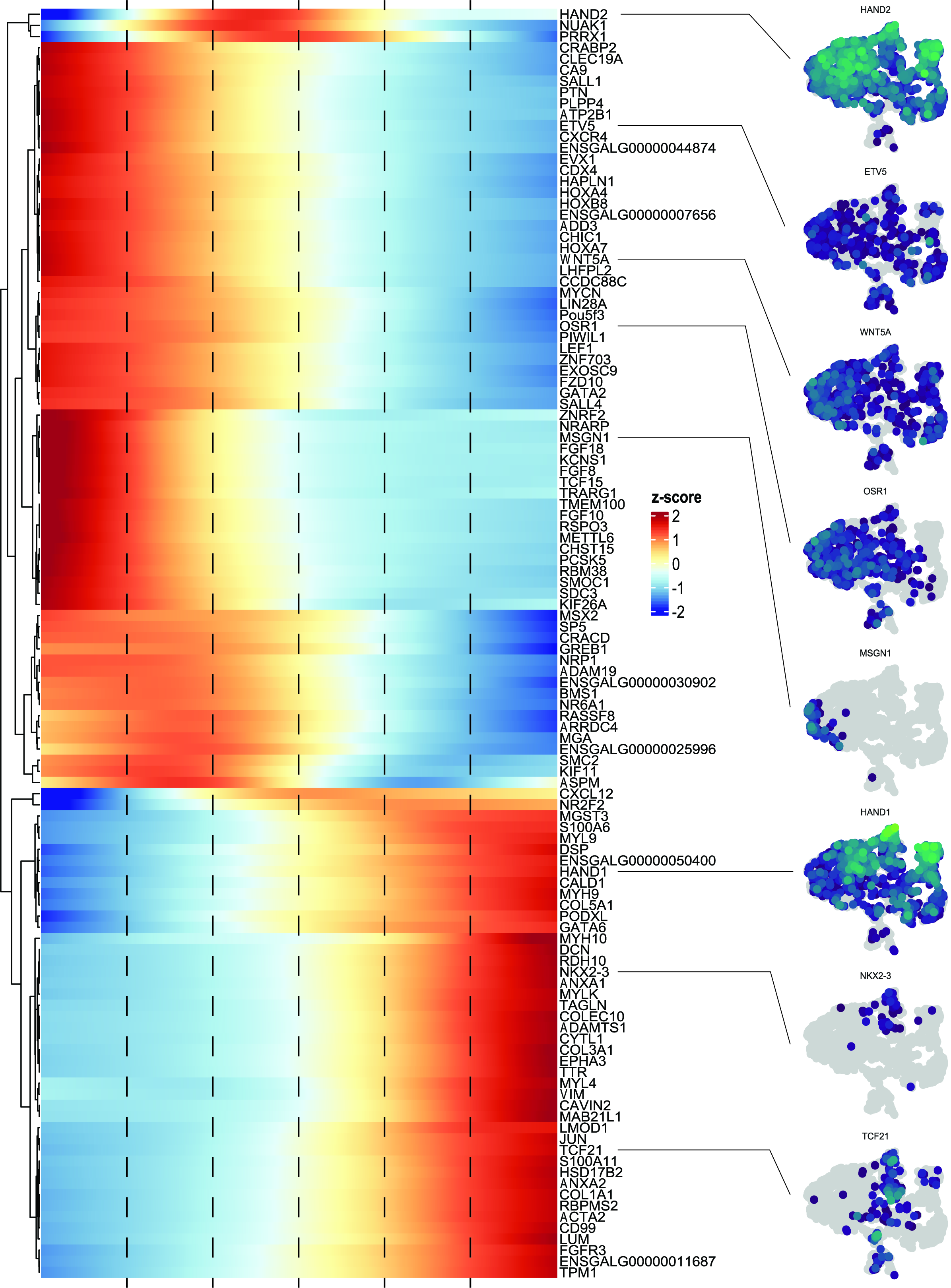

### Figure S3

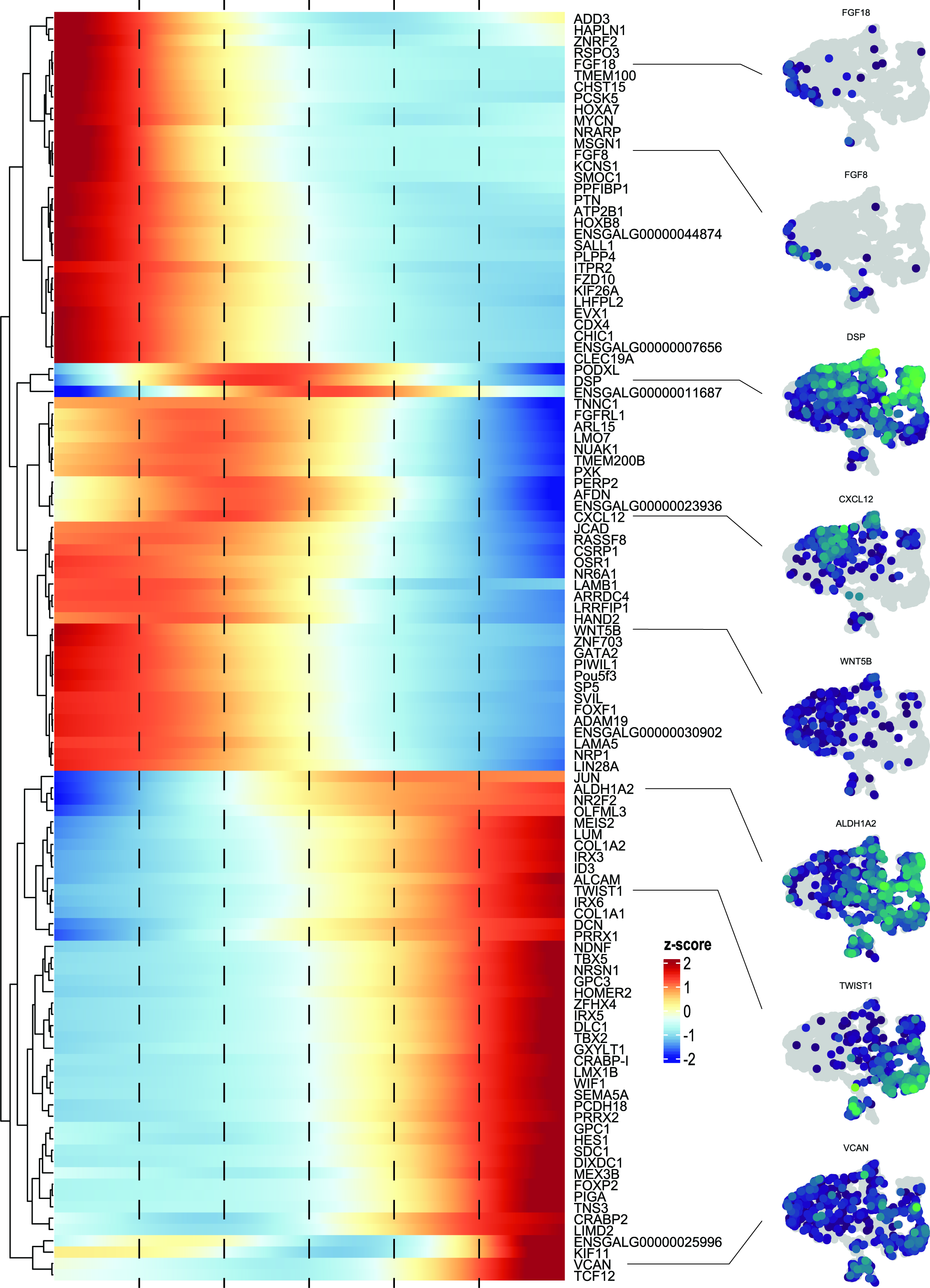

### Figure S4

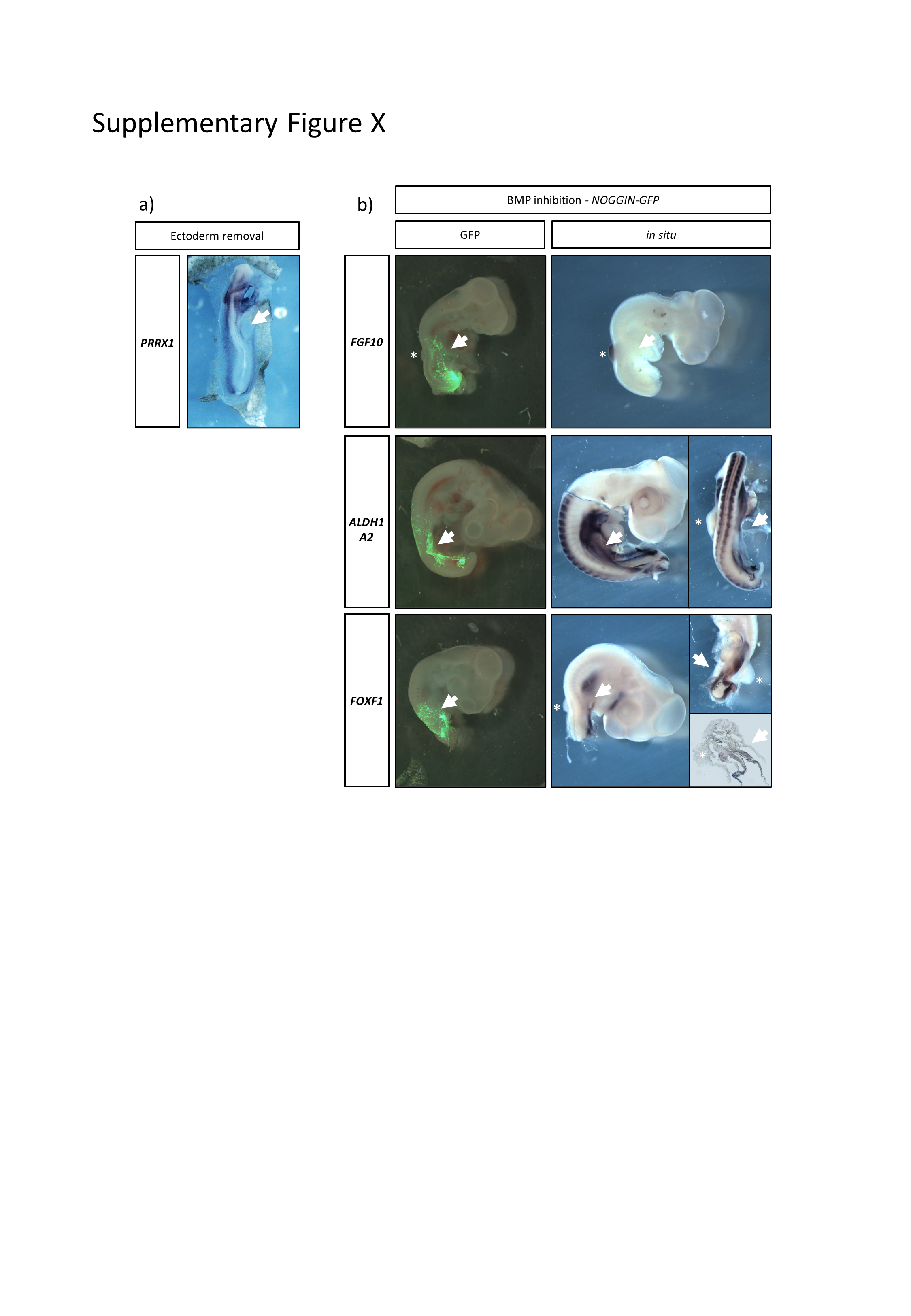
